# Compartmental Modeling Software: a fast, discrete stochastic framework for biochemical and epidemiological simulation

**DOI:** 10.1101/609172

**Authors:** Christopher W. Lorton, Joshua L. Proctor, Min K. Roh, Philip A. Welkhoff

**Author notes:** Co-first author.

## Abstract

The compartmental modeling software (CMS) is an open source computational framework that can simulate discrete, stochastic reaction models which are often utilized to describe complex systems from epidemiology and systems biology. In this article, we report the computational requirements, the novel input model language, the available numerical solvers, and the output file format for CMS. In addition, the CMS code repository also includes a library of example model files, unit and regression tests, and documentation. Two examples, one from systems biology and the other from computational epidemiology, are included that highlight the functionality of CMS. We believe the creation of computational frameworks such as CMS will advance our scientific understanding of complex systems as well as encourage collaborative efforts for code development and knowledge sharing.

## 1 Introduction

Developing fast, efficient, and scalable computational frameworks is integral to investigating a broad set of epidemiological and biological systems. Here, we present a new open-source computational framework, called the compartmental modeling software (CMS), which enables the simulation of discrete, stochastic reaction models. The CMS framework is highly flexible: the new model description language enables rapid model development; a broad set of available numerical algorithms allows users to optimize simulations based on model structure or computational speed requirements; and the standardized model output empowers a wide-variety a visualization tools. In this article, we report the functionality of the CMS framework while highlighting the key components of the software.

Open-source frameworks have been previously developed for discrete, stochastic compartmental modeling [20], including advancements that account for event handling [25], access via open-source software such as Python [6], and adaptations to cloud-based platforms [15]. Our new CMS framework broadens the scope and scale of previous open-source software. In addition to the standard exact and approximate numerical algorithms, CMS includes more recently developed rare-event probability estimation algorithms [13, 24]. Spatial simulation algorithms [22, 14, 9] are also integrated to accommodate spatial diffusion processes. Reaction propensities, time delays [8, 10], and state- and time-dependent events can be fully customized in the CMS framework. Coupling these features with software documentation, unit and regression tests, and objectoriented develop ment, CMS is a novel and flexible framework to allow for modeling of complex, physical systems.

**Fig. 1.**
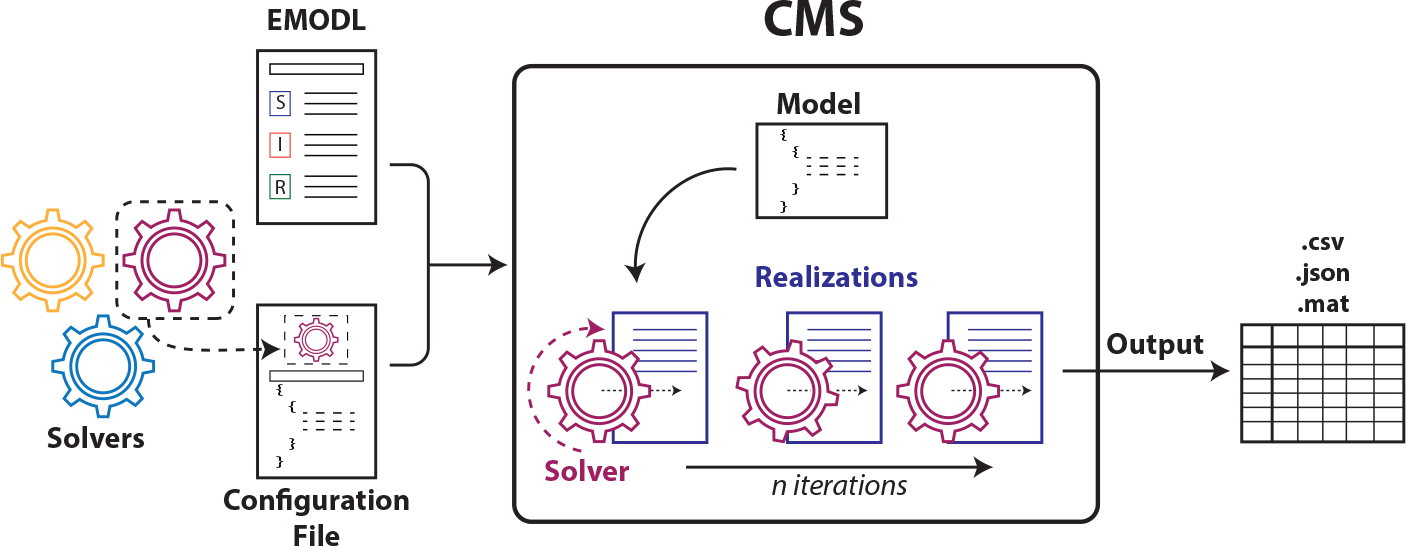
An overview of the CMS software.

## 2 Compartmental Modeling Software

In this section, we briefly explain the major components of CMS. An overview of the CMS schematics is given in Figure 1. For more detail, see the CMS documentation [1] as well as the GitHub repository for the associated code [2].

### 2.1 Platform and Computational Requirements

CMS was developed in C# and C++ and tested with NUnit framework [4]. It runs on 64-bit Windows 7 or later with any 64-bit Intel CPU. The CMS Visual Studio solution file has been updated to work with Visual Studio 2017 and targets the .NET Framework version 4.6.

### 2.2 Execution Pathways

Command line invocation is the most common usage for CMS. However, CMS can be executed in any language or scripting tool that can load Microsoft .NET technology. For example, CMS can be integrated into MATLAB or into Python through a package such as pythonnet [5].

### 2.3 Input Language and Configuration

A custom model description language, named the epidemiological modeling language (EMODL), was created to support the unique features of CMS and accommodate future expansion. EMODL syntax allows for efficient formulation of time- and state-based events as well as delays. It also supports custom propensity formulation in addition to the traditional propensities for mass-action kinetics. Runtime and solver-specific parameters are listed in a JavaScript Object Notation (JSON) [3] formatted configuration file. Parameters such as solver name, ensemble size, length of simulation, random number generator and output format are included in this configuration file.

### 2.4 Discrete Stochastic Solvers in CMS

CMS offers a comprehensive suite of stochastic solvers. It contains a total of 16 solvers, whose capabilities range from exact sampling of the true underlying probability distribution of the master equation (ME), fast approximation of the ME, spatial (network) simulations, and rare event probability estimation. We briefly describe each category below. In each subsection, we refer the reader to the original publication for mathematical derivations and computational complexity.

#### Exact Methods

The time evolution of systems are fully described by the solution of the ME, which provides the probability of every possible system state configuration at any given time. The ME is analytically intractable for most systems, but exact numerical solvers can estimate these probabilities numerically by sampling from the ME. Gillespie’s stochastic simulation algorithm (SSA) [18] is the most well-known exact method for simulating systems in stochastic chemical kinetics. CMS offers two known implementations of SSA—direct method and first reaction method—that are theoretically equivalent to each other. CMS also features Gibson and Bruck’s next reaction method [17] as well as the SSA with time delays [8, 10].

#### Approximate Methods

While the exact methods produce accurate time trajectories, explicitly simulating every reaction may be prohibitively slow for some systems. Approximate algorithms have been developed to accelerate the simulation at the expense of accuracy. CMS includes three approximate solvers—two implementations of *τ*-leaping [12] and one of R-leaping [7].

Tau-leaping is the most popularly used approximate method, and CMS offers two implementations for choosing a time step (*τ*)—adaptive and fixed. The adaptive time step selection mechanism also includes a check to avoid negative population [11] by reverting to SSA when appropriate. The fixed time step method assumes that the chosen *τ* is small enough to produce accurate trajectories. When this assumption is not met, negative population counts can occur.

#### Spatial Simulation Methods

Spatial simulation in CMS is possible via three different solvers—inhomogeneous SSA (ISSA) [22], diffusive finite state projection (DFSP) [14], and fractional diffusion (FD) [9]. ISSA divides a system into homogeneous subvolumes, and diffusive transfers are treated as a unimolecular reaction. Therefore, it can be prohibitively slow when fast diffusion is present. The other two methods are more efficient than the ISSA when diffusion occurs frequently with respect to the number of reaction events. DFSP solves the diffusion master equation by adapting the Finite State Projection (FSP) method [23], while FD is based on Lie-Trotter operator splitting of the diffusion and reaction terms. Unlike ISSA and DFSP, fractional diffusion allows for jumping to a distant locale with non-zero probability.

#### Rare Event Probability Methods

In addition to generation of time trajectories, CMS allows for efficient estimation of a rare event probability via the doubly weighted SSA (dwSSA) [13] and the state-dependent dwSSA (sdwSSA) [24]. Both algorithms utilize importance sampling, whose optimal parameters are determined by the information-theoretic concept of cross entropy. While the dwSSA assigns a single importance sampling parameter per reaction, the sdwSSA creates a list of state-dependent importance sampling parameters in order to further reduce the variance in the rare event probability estimate.

#### Exploratory Methods

The CMS framework is designed to enable efficient prototyping of new methods. A new solver can be easily implemented by extending the base solver. Four prototype methods are included in the CMS, and we refer to the documentation page [1] for further details. We note that solvers in this category are included as an example of ongoing method development and are not currently supported by the developers.

### 2.5 Output Files

There are three output formats available in the CMS—comma-separated values (CSV), JSON [3], and MATLAB (MAT). By default, CMS creates trajectories.csv in the output directory, with the realization index appended to each observable name specified in the EMODL file. Output-related options, such as compression and heading, can be specified in the configuration file.

## 3 Examples

### 3.1 Schlögl Process

The Schlögl process is a canonical example of a chemical system exhibiting bistability. This model consists of the following four reactions:

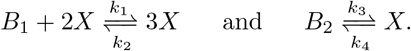

We take the system description including parameters and initial condition from [19] that produces bistable behavior in *X*: *k*_1_ = 3 × 10^−7^, *k*_2_ = 10^−4^, *k*_3_ = 10^−3^, *k*_4_ = 3.5, and x_0_ = [10^5^, 2 × 10^5^, 250], where **x**_0_ denotes the initial population of *B*_1_, *B*_2_, and *X*. Using the SSA solver, an ensemble of *N* = 1e5 simulations were generated to produce the distribution of *X* shown in Figure 2(a).

**Fig. 2.**
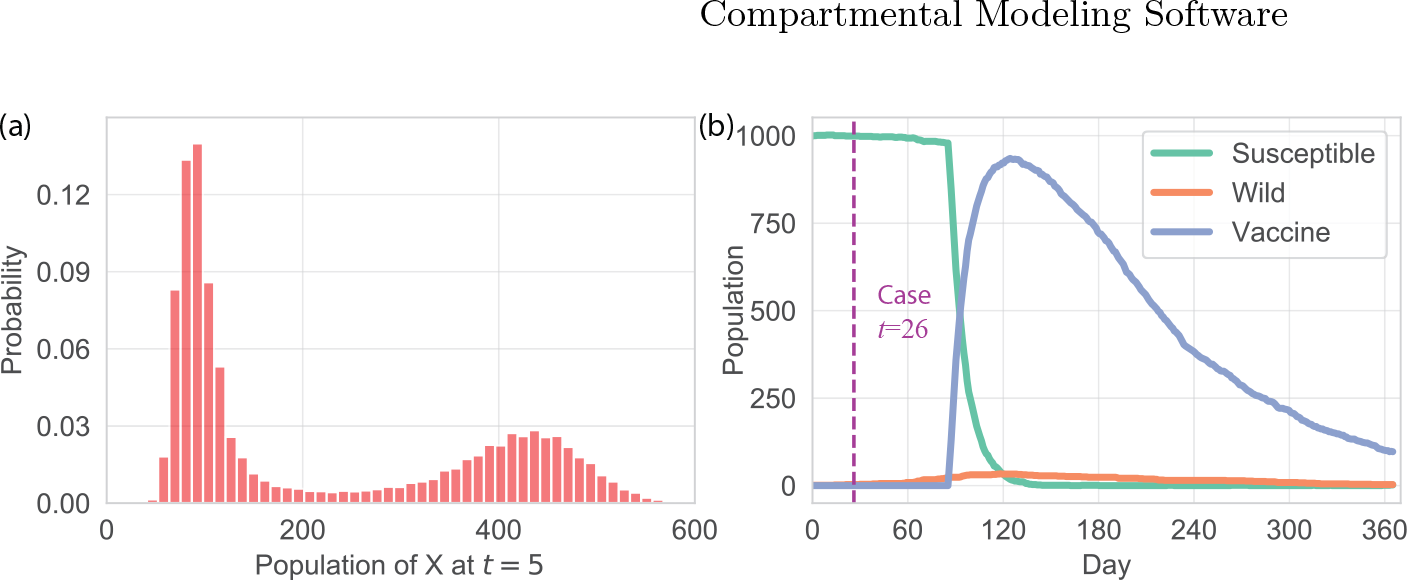
(a) illustrates the bistable distribution of population of species *X* at the final time *t* = 5. (b) illustrates one realization of a stochastic Polio outbreak, a paralyzed child (case) detection, and the delayed start of a vaccination campaign.

### 3.2 Vaccination campaigns for eradicating poliomyelitis

Globally, the number of poliomyelitis cases has dramatically decreased over the past two decades and may be the second human infectious disease to be eradicated [21]. The broad functionality of CMS can be utilized to model current vaccination questions such as programmatic mobilization time to arrest a small polio outbreak. We implement an established discrete, stochastic compartmental model of the spread of Polio [16]. We add two essential components to this model: a probabilistic case detection matching the 1 to 200 case to infection rate for polio and a vaccination campaign that is initiated after a fixed time duration following case detection.

We simulate the model starting with a population of 1000 individuals and one imported infection. When a case is detected, the model enforces a sixty day operational delay before a vaccination campaign begins. Figure 2(b) illustrates one realization of this model; note that a single case is detected on day 26 and vaccination begins on day 86. Due to the vaccination in this scenario, we do not detect another paralyzed child. To generate this realization, we utilize the SSA numerical algorithm, allow for a simulation duration of 365 days, and sample the state of the system each day. The model file, config file, and output file for this example can be found in the examples folder at [2].

## 4 Conclusion

Simulating biological and epidemiological processes with low population counts, such as nearing elimination of an infectious disease, requires the ability to simulate discrete, stochastic reaction models. The compartmental modeling software (CMS) is a novel, extensible framework used to generate an ensemble of trajectories that approximate the true underlying probability distribution described by the model and initial condition. CMS was designed to enable rapid model development with a custom model language, allow for user flexibility in choosing model-specific numerical solvers, and output the model trajectories into easy to visualize formats. Moreover, CMS is an open-source project allowing for community development; the object-oriented programming and class structure of the code allows for intuitive modifications of the code-base such as the inclusion of new numerical solvers or random number generators.

A number of challenges face the widespread adoption of our framework as a modeling tool. The source code has been written in C# which is not widely utilized in university settings. Also, CMS does not currently make use of multithreading, multi-core CPUs, or GPU resources. Despite these limitations, CMS can be deployed across multiple virtual machines since the memory and disk requirements are minimal. We also plan on providing a docker image for development and a reproducible environment for execution as well as developing an API from Python to enable users to call into the CMS executable. More broadly, we believe computational tools such as CMS will help provide insights into realistic biological and epidemiological systems.

## Acknowledgements

JLP, MKR, CWL, and PW would like to thank Bill and Melinda Gates for their active support of the Institute for Disease Modeling and their sponsorship through the Global Good Fund. The authors would also like to thank Mandy Izzo for her assistance illustrating Figure 1.

